# Progression of herpesvirus infection remodels mitochondrial organization and metabolism

**DOI:** 10.1101/2023.11.16.567337

**Authors:** Simon Leclerc, Alka Gupta, Visa Ruokolainen, Jian-Hua Chen, Kari Kunnas, Axel A. Ekman, Henri Niskanen, Ilya Belevich, Helena Vihinen, Paula Turkki, Ana J. Perez-Berna, Sergey Kapishnikov, Elina Mäntylä, Maria Harkiolaki, Eric Dufour, Vesa Hytönen, Eva Pereiro, Tony McEnroe, Kenneth Fahy, Minna U. Kaikkonen, Eija Jokitalo, Carolyn A. Larabell, Venera Weinhardt, Salla Mattola, Vesa Aho, Maija Vihinen-Ranta

## Abstract

Viruses target mitochondria to promote their replication, and infection-induced stress during the progression of infection leads to the regulation of antiviral defenses and mitochondrial metabolism which are opposed by counteracting viral factors. The precise structural and functional changes that underlie how mitochondria react to the infection remain largely unclear. Here we show extensive transcriptional remodeling of protein-encoding host genes involved in the respiratory chain, apoptosis, and structural organization of mitochondria as herpes simplex virus type 1 lytic infection proceeds from early to late stages of infection. High-resolution microscopy and interaction analyses unveiled infection-induced emergence of rough, thin, and elongated mitochondria relocalized at the perinuclear area, a significant increase in the number and clustering of ER-mitochondria contact sites, and thickening and shortening of mitochondrial cristae. Finally, metabolic analyses demonstrated that reactivation of ATP production is accompanied by increased mitochondrial Ca^2+^ content and proton leakage as the infection proceeds. Overall, the significant structural and functional changes in the mitochondria triggered by the viral invasion are tightly connected to the progression of the virus infection.

## Introduction

Mitochondria are double membrane-bound, ATP-producing powerhouses of the cells comprising approximately 1,000 proteins^1^, and mitochondrial metabolism produces precursors for biosynthetic pathways including nucleotides, lipids, and amino acids^2–4^. The mitochondria also have a role in the innate immune system including the ability to respond to various types of internal and external stressors. Both the internal stressors such as genetic, metabolic, and biochemical factors, as well as external stressors such as environmental agents lead to dynamic functional and morphological alterations of the mitochondria^5,6^.

One of the external stressors that affect the mitochondria is a viral infection^7^. Production of viral proteins is followed by stimulation of mitochondria-mediated immune signaling and cell death pathways^8,9^. The mitochondria have an important role in the mediation of programmed cell death, and viruses target this machinery either to ensure the viability of cells for the viral replication or to destroy them for more efficient spreading of progeny viruses^10^. Many mitochondrial changes, including those related to apoptosis, are triggered by calcium influx from the endoplasmic reticulum. The mitochondria and ER are often physically located next to each other and connected via specific contact sites, and multiple tethering complexes between mitochondria and the ER have been identified, including VAPB-PTPIP51^11^. Several viruses have been shown to target the calcium homeostasis between the ER and mitochondria either to promote or inhibit apoptosis^12–14^. Besides using various mechanisms to evade cellular antiviral responses, the progression of viral replication relies on the reprogramming of the host cell mitochondrial metabolism. Severe acute respiratory syndrome coronavirus 2 (SARS-CoV-2) and human immunodeficiency virus (HIV) infections induce disturbance of mitochondrial homeostasis, which leads to increased glycolysis and production of metabolites useful in the synthesis of lipids, nucleotides, and proteins needed during the formation of viral particles^15–18^. Herpes simplex virus type 1 (HSV-1) supports the host cell mitochondrial oxidative and biosynthetic metabolism by inducing replenishment of tricarboxylic acid cycle intermediates^19^. Another herpesvirus, human cytomegalovirus (HCMV), activates mitochondrial energy production by increasing glycolysis^19,20^.

The alpha-herpesvirus HSV-1 is an enveloped double-strand DNA virus that hijacks the cellular metabolism including mitochondrial biosynthetic pathways to produce precursors required for replication^21^. The regulation of apoptosis to prevent the premature death of the host cells is essential for the envelopment and nonlytic cellular egress of HSV-1^9,22,23^. The viral proteins ICP4 and Us3 suppress apoptosis by inhibiting caspase activation and inactivating proapoptotic proteins, and ICP27 protein counteracts caspase 1-dependent cell death^24–26^. The HSV-1 infection has been shown to cause relocalization of mitochondria towards the nuclei as well as elongation of mitochondria^27,28^. In the absence of infection, elongation can be induced by starvation and inflammatory response^29^. The elongation can protect mitochondria from autophagy and lead to an increase in the surface area of cristae to enhance ATP production^30^.

Here, we analyze how the progression of HSV-1 infection influences mitochondrial gene transcription, ultrastructure, and organization, and how it contributes to mitochondrial function. We use various microscopical and biochemical tools uniquely suited for specific observing of detailed changes in mitochondrial organization and function of HSV-1-infected cells. Our approach provides characterization and quantification metrics for the mitochondrial protein transcription, mitochondrial structures including cristae, and energy metabolism at the early and late stages of infection. We demonstrate that late infection-induced significant changes in the mitochondrial protein gene transcription are associated with dramatic changes in the mitochondrial ultrastructure and distribution, ER contacts, and function. Our results illustrate how a pathogenic DNA-virus infection remodels the structural organization and function of the mitochondria during the progression of the lytic infection.

## Results

### The transcription of mitochondria-associated proteins is modified in HSV-1 infection

To explore virus-induced changes in the host cell transcription, we employed a global run-on sequencing (GRO-seq) analysis of nuclear-encoded genes in green monkey kidney epithelial (Vero) cells. The clustering of genes in the same pathways (based on the STRING database (https://string-db.org/) of predicted functional protein-protein interactions) revealed that the infection-induced changes in the gene transcription were linked to proteins that contribute to mitochondrial functions such as electron transport, immunity, and apoptosis (Fig. 1a,b). A substantial increase in functional interactions and regulation of transcription was detected when infection proceeded from 4 to 8 hpi. (Fig. 1a,b, Supplementary Fig. 1a,b). In general, the transcription of 43 and 59 nuclear-encoded mitochondrial genes (hereinafter simply referred to as mitochondrial genes) was upregulated, while 22 and 102 were downregulated at 4 and 8 hpi, respectively, in comparison to the noninfected cells (Supplementary Fig. 1b). Overall, the progression of infection reduced the transcription of genes involved in energy metabolism. One of the key components of the electron transport chain, the first enzyme of the respiratory chain, is complex I (NADH-ubiquinone oxidoreductase)^31^. Two and nine of the genes encoding proteins involved in this 45-protein complex, consisting mostly of nuclear-encoded mitochondrial proteins (NDUF subunits), were downregulated at 4 and 8 hpi, respectively. As expected, activation of mitochondrial antiviral defenses led to the regulation of genes associated with apoptosis and immune response (Fig. 1a,b, Supplementary Fig. 1b). For example, the apoptosis-inducing protein BCL2L11 (also known as BIM), a member of the BCL2 family regulating the intrinsic, mitochondrion-dependent apoptosis^32^, was upregulated both at 4 and 8 hpi. Transcription of the gene encoding histone deacetylase Sirtuin 1 (Sirt1), a multifunctional protein regulating apoptosis by controlling the cellular distribution of p53^33^, also increased in infected cells (Fig. 1a,b, Supplementary Table 1). In contrast, transcription of the gene encoding tumor necrosis factor receptor-associated factor 6 (TRAF6), a member of the TRAF family of proteins mediating signaling pathways regulating inflammatory signaling, was downregulated both at 4 and 8 hpi. By suppressing cell death complex assembly, TRAF6 can inhibit tumor necrosis factor α (TNF-α)-induced apoptosis and necroptosis^34^. Moreover, the upregulated transcription of ATAD3A of membrane protein AAA domain-containing protein 3 member A (ATAD3A) at 4 hpi, involved in the removal of damaged mtDNA^35^, is in line with the HSV-1 infection-induced disintegration of the mtDNA^27,36^. Finally, the infection increased the transcription of mitochondrial calcium voltage-gated channel subunit alpha1 B (CACNA1B) at 4 and 8 hpi.

**Figure 1.**
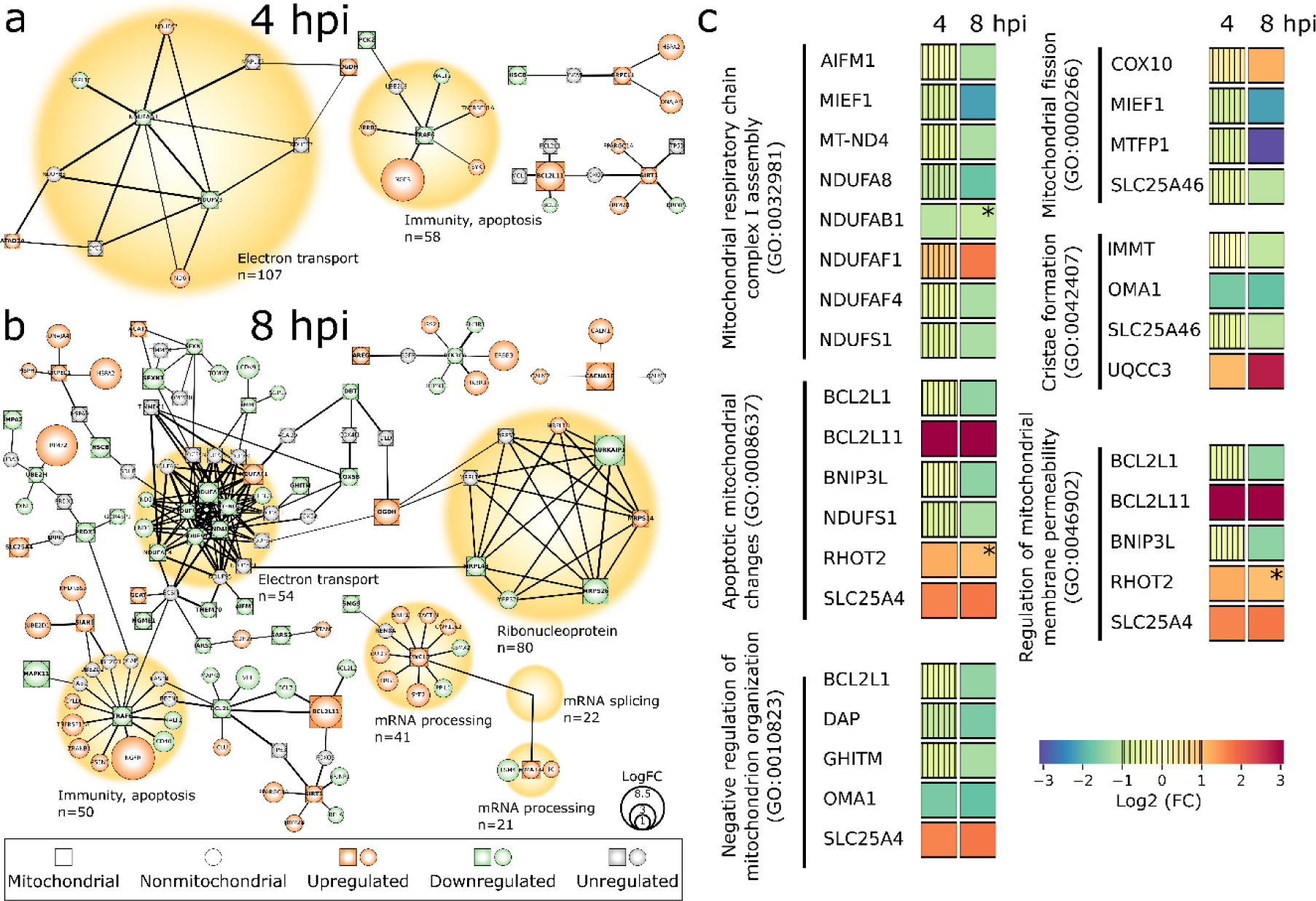
HSV-1 infection alters the host transcriptome. Global run-on sequencing (GRO-seq) analysis of nascent RNA levels of mitochondrial and mitochondria-associated proteins in infected Vero cells at **(a)** 4 and **(b)** 8 hpi. The genes encoding for mitochondrial proteins (square-shaped nodes, bolded titles) and nonmitochondrial cellular interactor proteins (round-shaped nodes) are shown. The upregulated (orange) or downregulated (green) transcripts in response to infection are visible together with unregulated interacting transcripts (grey). The node size correlates with a logarithmic fold change (logFC) of regulation and the thickness of black lines between nodes is proportional to the interaction in the STRING database (https://string-db.org/). The main functions of the proteins are denoted with yellow circles with their size proportional to the number of interactors. **(c)** GO term classification for mitochondrial processes in the infected cells at 4 and 8 hpi. The color bar indicates upregulation (orange-red) or downregulation (green-blue), and low change of gene transcription (vertical stripes). The non-significant enrichment is also shown (*).

To further assess the role of mitochondrial genes differentially expressed during the infection, we performed the gene ontology analysis (GO) according to the PANTHER classification system for the GO Biological process (http://www.pantherdb.org). The enriched GO terms associated with the regulation of mitochondrial organization and function were mitochondrial respiratory chain complex I assembly (GO:0032981), apoptotic mitochondrial changes (GO:0008637), negative regulation of mitochondrion organization (GO:0010823), mitochondrial fission (GO:0000266), cristae formation (GO:0042407), and regulation of mitochondrial membrane permeability (GO:0046902) (Fig. 1c, Supplementary Table 2). The transcription of genes encoding proteins in the mitochondrial respiratory chain was mostly downregulated at 8 hpi. The downregulated genes corresponding to complex I proteins included NADH dehydrogenase subunit 4 (MT-ND4) and several nuclear-encoded NDUF subunits, and only the gene encoding NDUFAF1 was upregulated. In general, infection-induced stress results in cellular and viral pro- and antiapoptotic responses. Viral factors aim to enhance the envelopment and cellular exit of HSV-1 virions by preventing premature cell death^37,38^. On the other hand, we found that cellular proapoptotic processes included the upregulation of gene encoding apoptosis-inducing protein BCL2L11 and an integral outer mitochondrial membrane protein SLC25A46 (also known as adenine nucleotide translocator 1, ANT1). SLC25A46 is a multifunctional protein involved in mitochondrial ultrastructure alteration including the formation of elongated mitochondria, cristae maintenance, the enhancement of mitochondrial respiration/glycolysis, reactive oxygen species production, oxidative stress, and function of mitochondrial/ER contact that facilitates lipid transfer^39,40^. In infected cells, the upregulation of SLC25A46 was also involved in the negative regulation of mitochondrion organization and regulation of mitochondrial membrane permeability affecting proton gradient (Fig. 1c, Supplementary Table 2). Moreover, the gene encoding ubiquinol-cytochrome C reductase complex assembly factor 3 (UQCC3 also known as C11orf83), with a role in cristae formation^41^, was upregulated at 8 hpi. The downregulation of mitochondrial inner membrane protease (OMA1) is involved in the negative regulation of mitochondrion organization and cristae formation^42^. Activation of OMA1 during apoptosis and cellular stress leads to cleavage of another membrane protein, optical nerve atrophy 1 (OPA1), and results in the remodeling of cristae, the release of cytochrome c, and fragmentation of mitochondria^42,43^. Also, the genes encoding growth hormone-inducible transmembrane protein (GHITM, also known as TMBIM5 or MICS1), mitochondrial fission process 1 (MTFP1), and the inner membrane mitochondrial protein (IMMT, also known as Mic60, HMP), with roles in mitochondria morphology dynamics, fission, cristae organization, and contact sites^44–47^, were downregulated at 8 hpi.

Altogether, we conclude that the progression of infection triggers timely responses of mitochondrial functions through regulation of transcription. The up- and downregulation of nuclear-encoded mitochondrial genes most likely reflect the balance between the cellular counteracts against the viral infection and the viral attempts to change cell functions to favor viral replication.

### Infection triggers changes in mitochondrial size and shape

Progression Soft X-ray imaging is a non-invasive method for high-contrast imaging of the internal structure of intact frozen cells^48^. This method allows label-free imaging of carbon- and nitrogen-containing sub-cellular organelles. The image contrast arises from the attenuation of soft X-rays when they interact with the cellular internal structures. Specifically, densely packed proteins or lipids attenuate strongly^49–51^. To create a comprehensive view of virus-induced changes in the 3D structure and distribution of mitochondria, we analyzed suspended adherent mouse embryonic fibroblast (MEF) cells in glass capillaries and adherent MEF cells grown on EM grids using cryo soft X-ray tomography (SXT). Cryo-SXT images of suspended cells showed the development of elongated mitochondria at 4 hpi and increasingly at 8 hpi (Fig. 2a, b, Supplementary movie 1). The linear absorption coefficient (LAC) values from the isotropic reconstructed 3D tomograms, depending on the concentration and composition of biomolecules such as lipids and proteins, were measured for each voxel of the segmented 3D mitochondria. The LAC values showed a significant increase at 4 and 8 hpi compared to the noninfected cells. This suggests that the infection led to an increased density of mitochondrial biomolecules (Fig. 2c). Cryo-SXT data analysis also showed that the number of mitochondria increased at 4 hpi, and then decreased as the infection progressed to 8 hpi, suggesting that the fragmentation of mitochondria at early infection was followed by their fusion at late stages of infection (Fig. 2d). However, the average volume of mitochondria increased as infection proceeded which argues against the disintegration (Fig. 2e, Supplementary Fig. 2c). As the segmentation of SXT data was challenged by artificial fragmentation of elongated mitochondria, we decided to apply serial block face scanning electron microscopy (SBF-SEM) data, which is better suited for 3D analysis of mitochondrial length. We found that the length of mitochondria increased at 8 hpi (2.8 ± 0.4 µm) and at 12 hpi (2.9 ± 0.2 µm) in comparison to noninfected cells (2.0 ± 0.1 µm) (Supplementary Fig. 2 a,b). Altogether, this suggests that the progression of infection triggers mitochondrial elongation accompanied by an increase in the volume. The cryo-SXT imaging of adherent cells confirmed the infection time-dependent elongation of mitochondria (Fig. 2f), and 3D reconstructions of the data revealed increased roughness of the mitochondrial outer surface at 8 and 12 hpi (Fig. 2g, Supplementary Movie 2). By 12 hpi, the mitochondria had moved closer to the nuclear envelope (Fig. 2h) and they became thinner (Fig. 2i). Finally, the analysis of the ratio between the mitochondrial surface area and volume confirmed an increase in the surface roughness at late infection (Fig. 2j). Taken together, the progression of infection resulted in the increased emergence of elongated and thin mitochondria with a rougher surface that accumulated closer to the nuclear envelope.

**Figure 2.**
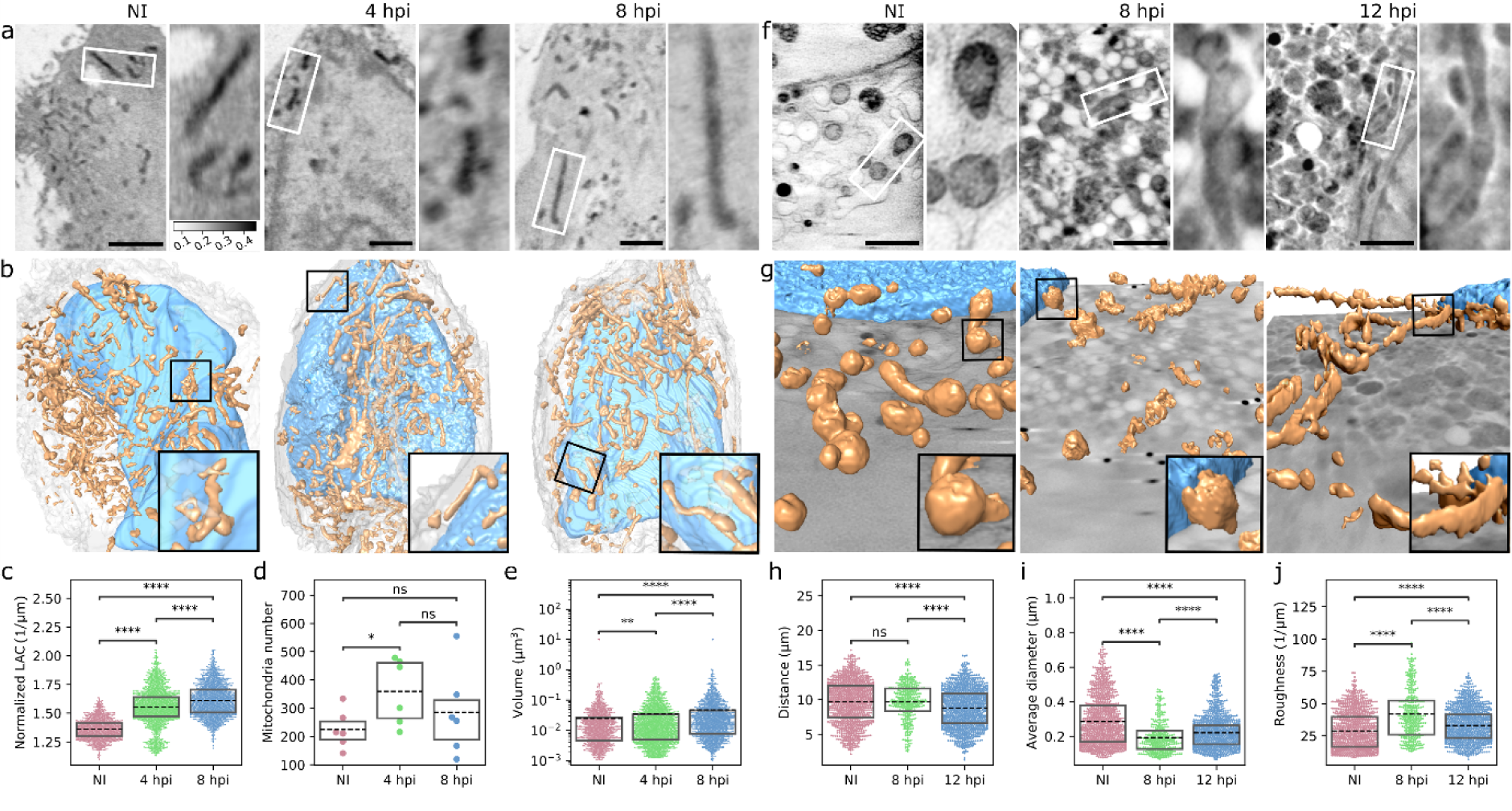
The progression of infection leads to thiner and rougher mitochondria. **(a)** Cryo-soft X-ray tomography (SXT) thin slices (ortho slices) of suspended MEF cells in glass capillaries and **(b)** 3D reconstructions of segmented mitochondria (brown) in the cytoplasm around the nucleus (blue) in noninfected (NI) and infected MEF cells at 4 and 8 hpi. White and black squares show the magnified mitochondria. Scale bars, 2 µm. See also the Supplementary Movie 1. Box plots of (**c**) linear absorption coefficient (LAC) values (grayscale bar 0.1-0.5 1/microns), (**d**) number, and (**e**) volume of mitochondria in noninfected and infected cells (n_cells_ = 6, n_mito_ = 1349, 2158, and 1713 for NI, 4 and 8 hpi, respectively). (**f)** SXT images of adherent MEF cells grown on EM grids and (**g**) 3D presentations of segmented mitochondria (brown) and nuclear envelope (blue) in noninfected and infected MEF cells at 8, and 12 hpi. The magnified mitochondria are shown in white and black squares. Scale bars, 2 µm. See also the Supplementary Movie 2. (**h)** The distance of mitochondria from the nucleus, (**i**) the diameter along the short axis, and (**j**) surface-to-volume ratio analysis of mitochondrial roughness in noninfected and infected cells (n_cells_ = 12, 4, and 13, n_mito_ = 1089, 364, and 1165 for NI, 8, and 12 hpi, respectively). The box plots show the mean (dashed line) and the interquartile range. Statistical significance was determined using the Brunner-Munzel test. The significance values shown are denoted as **** (p<0.0001), * (p<0.05), or ns (not significant).

### ER-mitochondria interplay is increased in infection

The 10–30 nm distance between the ER and mitochondria allows the protein-mediated tethering of the ER and the outer mitochondria membrane (OMM)^52^. Multiple functions that occur at those contact sites include Ca^2+^ signaling, lipid biosynthesis, and mitochondrial division^53^. To observe in high resolution the infection-induced changes in the 3D ultrastructure and specifically in the regions of close contact between the ER and mitochondrial membranes, we used serial block face scanning electron microscopy (SBF-SEM) (Fig. 3a). The 3D reconstruction and analyses of the segmented ER and mitochondria from the SBF-SEM data revealed both smaller and more extensive contact sites (Fig. 3b, Supplementary Movie 3). Infection-induced changes led to a clear decrease in the distance between the ER and mitochondrial membranes at 8 hpi in comparison to noninfected control cells. At 12 hpi the distance increased in comparison to 8 hpi, showing that ER and mitochondria move further away from each other in the late stages of infection (Fig. 3c). This was supported by an increased number of contact sites and contact region area at 8 hpi and a slight decrease at 12 hpi (Fig. 3d,e).To study the ER-mitochondria linkage further, we analyzed the distribution of tethering protein located on the ER, namely vesicle-associated membrane protein B (VAPB), by using the ten-fold expansion microscopy^54^. In infected cells at 8 hpi VAPB was distributed heterogeneously and formed clusters along the mitochondria, in contrast to the more uniform distribution seen in noninfected cells (Fig. 4a). VAPB also moved further away from the mitochondria in infected cells when compared to noninfected cells (Fig. 4b). Having shown a change in VAPB distribution, we next analyzed ER-mitochondria contact sites by closeness of ER-mitochondria tethering proteins VAPB and regulator of microtubule dynamics 3 (RMDN3, also known as PTPIP51) using proximity ligation assay (PLA). PLA is an immunodetection technique that generates a fluorescent signal only when two antibodies against the antigens of interest are closer than 40 nm from each other^55^. PLA between VAPB and RMDN3 antibodies revealed an increased number of punctate PLA signals as the infection proceeded (Fig. 4c,d). Notably, the progression of infection also resulted in an increase in the volume of the PLA foci accompanied by an increased size variation (Fig. 4e). The presence of enlarged foci suggests that part of the contact sites form discrete clusters on the mitochondrial membrane as the infection proceeds. Overall, these analyses demonstrated that the progression of infection results in increased VAPB-RMDN3 tethering and clustering of ER-mitochondria contact sites.

**Figure 3.**
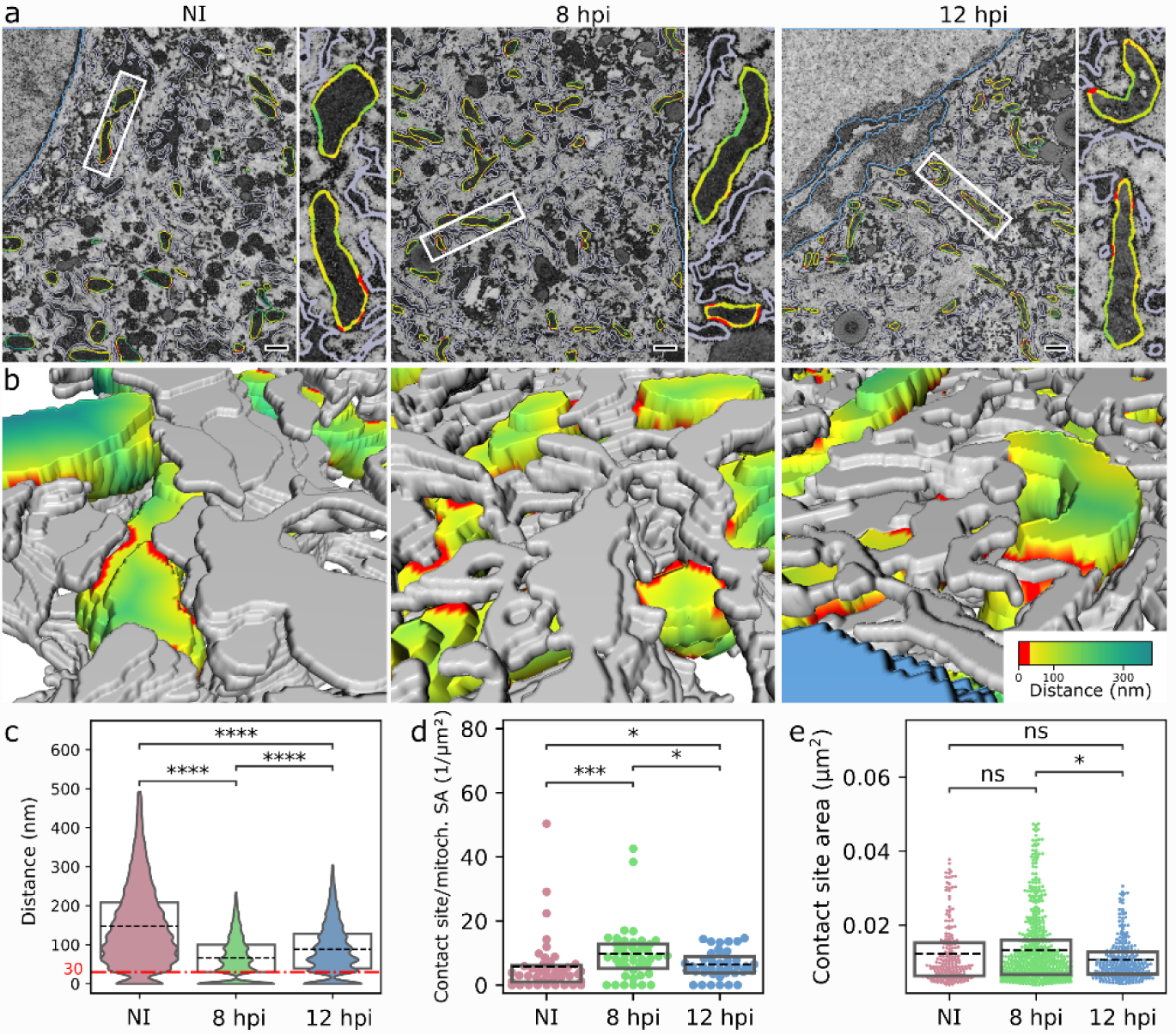
ER-mitochondria contacts increase in infection. **(a)** Representative sections of serial block face scanning electron microscopy (SBF-SEM) of noninfected and infected MEF cells at 8 and 12 hpi. White squares show the magnified mitochondria. The pseudocolor lines around the mitochondria show the closeness of ER and mitochondria, and regions of contact are defined as points where opposed membranes are within 30 nm of each other (red). Scale bars, 0.5 µm. (**b)** A higher-magnification 3D SBF-SEM reconstructions show the regions of contacts (distance less than 30 nm, red) between mitochondria (pseudocolor) and ER (grey). The pseudocolor bar indicates the distance between the ER and mitochondria. See also the Supplementary Movie 3. Violin and box plots show (**c**) the ER mitochondria distance, (**d**) the number (Nb) of contact sites/mitochondrial surface area, and **(e)** the area of the contact sites (n_mito_ = 43, 43, and 39 for NI, 8, 12 hpi, respectively). The box plots show the mean (dashed line) and the interquartile range. Statistical significance was determined using the Brunner-Munzel test. The significance values are denoted as **** (p<0.0001), *** (p<0.001), *(p<0.05), or ns (not significant).

**Figure 4.**
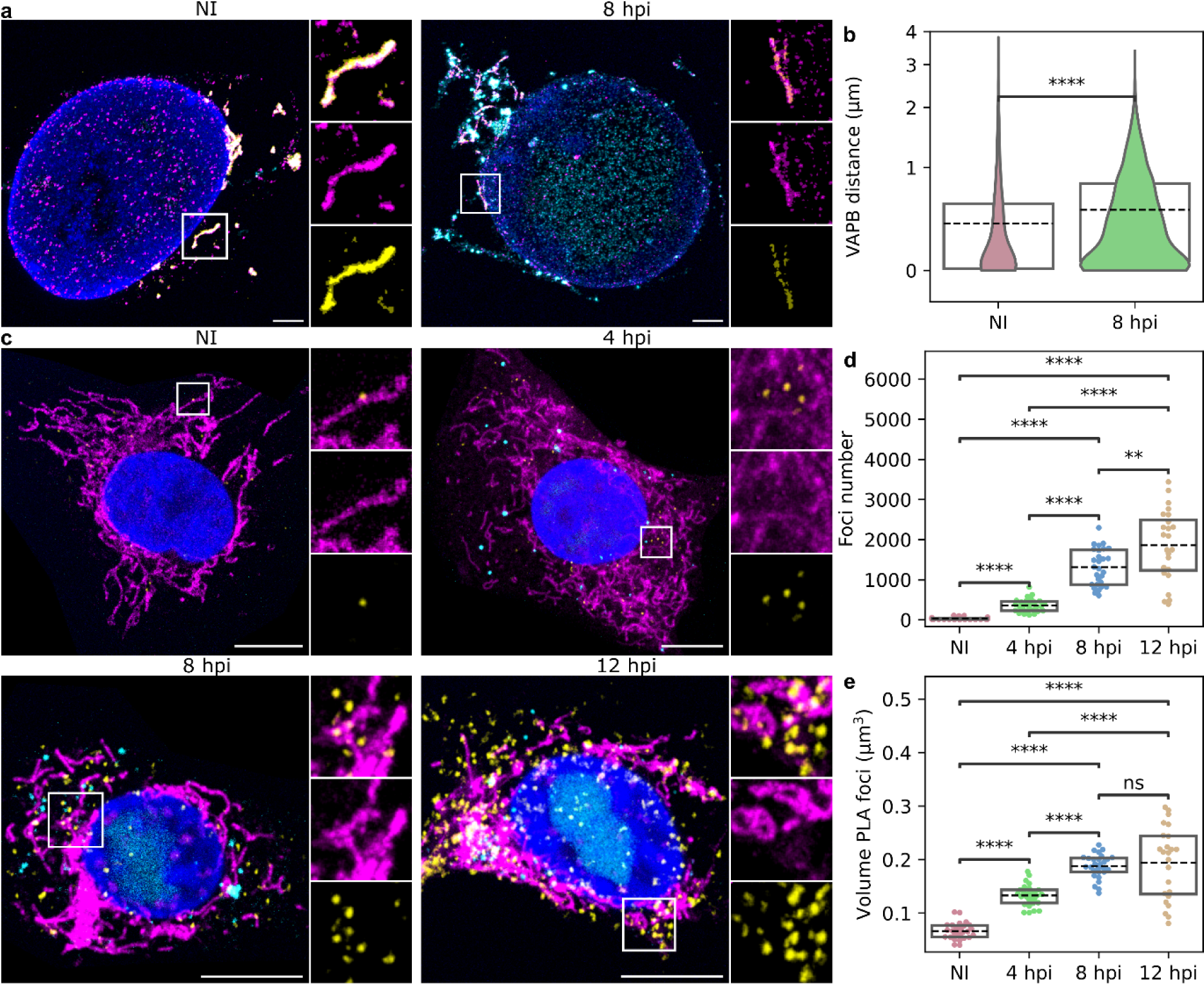
The amounts of ER-mitochondria contact sites increase and they are clustered in infection. **(a)** Visualization of the mitochondrial structure by the ten-fold robust expansion microscopy (TREx) in noninfected and infected Vero cells at 4, 8, and 12 hpi (n = 6). The distributions of an ER protein tyrosine vesicle-associated membrane protein B (VAPB, green) located in the ER-mitochondria contact sites and mitochondria labeled with MitoTracker (red) are shown. The localization of the viral replication compartment is presented by HSV-1 EYFP-ICP4 (cyan) and the nucleus by the DAPI stain (blue). Scale bars, 1 µm. (**b**) Violin plots show the distance between the mitochondria and VAPB. The red dashed line indicates the resolution threshold. (**c**) Proximity ligation analysis (PLA) of contact sites in noninfected and infected Vero cells at 4, 8, and 12 hpi. The PLA signal between VAPB and regulator of microtubule dynamics (RMDN3) is visualized by fluorescent spots (yellow). Mitochondria are labeled with MitoTracker (red), the nucleus with DAPI, and viral replication compartments are visualized by EYFP-ICP4. Scale bars, 10 µm. (**d**) Box plots showing the number of PLA foci and (**e**) volume of the foci per cell (n_cells_ = 27, 27, 28, and 24 for NI, 4, 8, and 12 hpi, respectively). The box plots show the mean (dashed line) and the interquartile range. Statistical significance was determined using the Student’s t-test. The significance values are denoted as **** (p<0.0001), ** (p<0.01), * (p<0.05) or ns (not significant).

### Mitochondrial cristae thicken and shorten as infection proceeds

Our GRO-seq findings demonstrating the upregulation of positive regulators and downregulation of negative regulators of cristae formation (Fig. 1c) suggested that the progression of infection leads to changes in cristae. We used focused ion-beam scanning electron microscopy (FIB-SEM) to characterize cristae 3D phenotypes as HSV-1 infection progresses in MEF cells (Fig. 5a). The mitochondria showed hallmarks of cristae remodeling at 8 hpi, including thickening and shortening of cristae (Fig. 5b, Supplementary Movie 4). Analyses of segmented and reconstructed image data showed that the average thickness of the lamella-shaped cristae increased at 8 hpi when compared to control cells and, specifically, some of the cristae were visibly swollen. In general, the thickness of cristae was more variable in infected than in noninfected cells (Fig. 5a, 5b, 5c). In comparison to noninfected cells, the progress of infection significantly shortened cristae (Fig. 5d) without changing their total surface area per mitochondrion (Fig. 5e).

**Figure 5.**
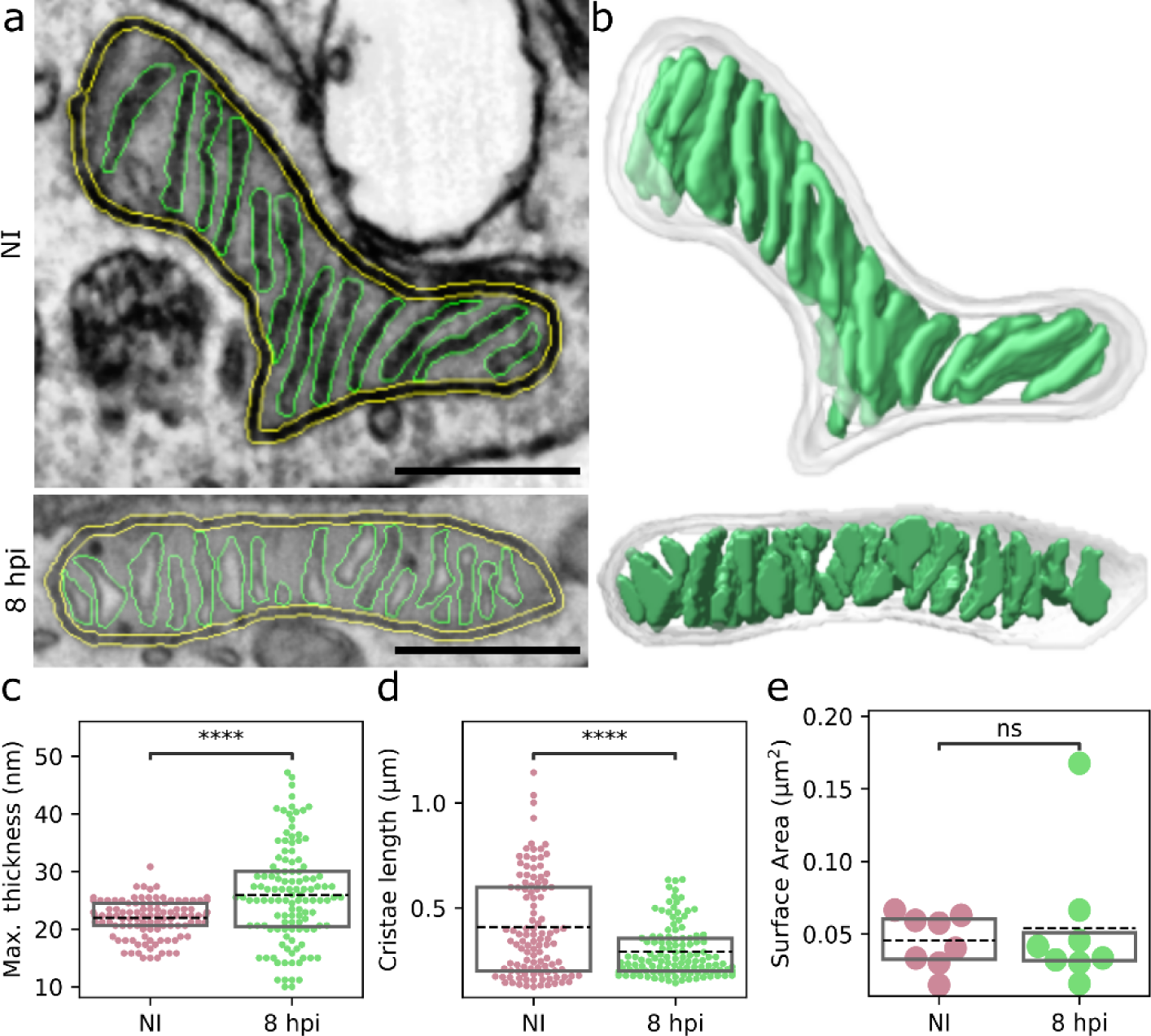
The cristae become thicker and shorter along the progression of the infection. **(a)** Representative focused ion-beam scanning electron microscopy (FIB-SEM) images of noninfected and HSV-1-infected MEF cells at 8 hpi. The mitochondrial outer membrane (yellow) and cristae (green) are shown. Scale bars, 0.5 µm. (**b**) The 3D structure of cristae (green) reconstructed from FIB-SEM stacks. See also the Supplementary Movie 4. The 3D quantitative analysis of (**c**) the maximal thickness and (**d**) the length of cristae calculated using a watershed algorithm to individualize the cristae lamella in noninfected and infected cells (n_mito_ = 8, n_crist_ = 112 and 121 for NI and 8 hpi, respectively). (**e**) The surface area of segmented cristae in each mitochondria. The box plots show the mean (dashed line) and the interquartile range. Statistical significance was determined using the Student’s t-test. The significance values are denoted as **** (p<0.0001) or ns (not significant).

### Infection increases mitochondrial proton leakage and Ca^2+^ content

We next asked whether the remodeling of cristae is associated with the modulation of mitochondrial energy metabolism. Oxygen consumption rate (OCR) of the host MEF cells was analyzed using the Seahorse XF24 analyzer (Fig. 6a). During the infection, basal respiration and ATP production decreased at 4 hpi and restored to the level of noninfected cells at 8 hpi (Fig. 6b,c). In contrast, the mitochondrial proton leakage into the matrix through the inner mitochondrial membrane significantly increased only at 8 hpi (Fig. 6d). The coupling efficiency, which compares how oxygen is distributed between ATP synthesis and proton leak, demonstrated that ATP synthesis efficiency was best in noninfected cells and decreased towards 8 hpi (Fig. 6e). Finally, single-cell fluorescence microscopy measurements revealed that the mitochondrial calcium uptake (Fig. 6f) increased at 4 hpi and even more at 8 hpi (Fig. 6g), which is consistent with the increase in the number of ER-mitochondria contact sites (Fig. 3) mediating Ca^2+^ import to mitochondria from the ER^56^. Altogether, ATP production, proton leakage, and mitochondrial [Ca^2+^] were more pronounced at 8 compared to 4 hpi suggesting a progressive change as a result of the progression of virus infection.

**Figure 6.**
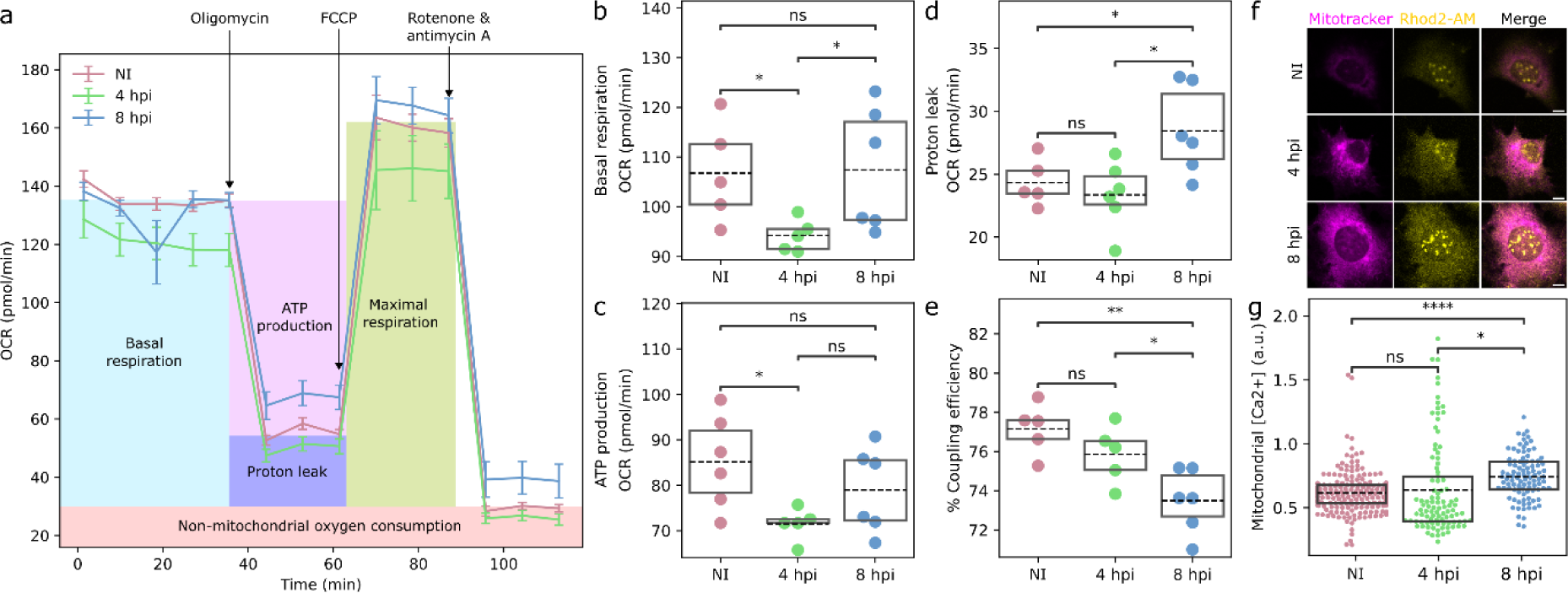
The infection leads to an increase in the mitochondrial proton leakage and Ca^2+^ content. **(a)** Seahorse real-time cell metabolic analysis of oxygen consumption rate (OCR) traces of noninfected and infected MEF cells at 4 and 8 hpi. The arrows mark the addition of oligomycin, FCCP, and rotenone + antimycin A (n = 30 000 cells, 6 replicates). Box plots showing (**b**) basal OCR, (**c**) proton leak (**c**) ATP production, and (**e**) coupling efficiency (ATP/O). (**f**) The live cell analysis of the mitochondrial calcium labelled by calcium ion indicator Rhod2-AM (green) selectively accumulating within mitochondria, and mitotracker (red). The infected cells selected for imaging were identified by the expression of HSV-1 ICP4 (not shown). Scale bars, 10 µm. (g) The quantitive analysis of mitochondrial [Ca^2+^] in noninfected and infected cells at 4 and 8 hpi (n = 207, 97, and 96 for NI, 4, and 8 hpi, respectively). The box plots show the mean (dashed line) and the interquartile range. Statistical significance was determined using the Student’s t-test. The significance values shown are denoted as **** (p<0.0001), ** (p<0.01), *(p<0.05), or ns (not significant).

## Discussion

In infected cells, there is a dynamic balance between viral reprogramming of cellular machinery to advance viral replication and counteracting the infection by cellular defenses. Mitochondrial metabolism and antiviral functions have a central role in this process. Our results demonstrate that the progression of HSV-1 infection results in extensive remodeling of mitochondrial gene transcription, organization, interactions, and energy metabolism.

During the early HSV-1 infection, energy is required for the stepwise progression of viral gene expression, genome replication, and assembly of progeny virions^21,57,58^. We show that the progression of HSV-1 infection from early to later phases results in a reduction in the transcription of genes encoding the proteins of the mitochondrial respiratory chain, specifically the proteins of complex I. Consistent with earlier studies, we show that the mitochondria are reassembled and mitochondrial ATP machinery is reactivated later and remains active until 12 hpi^27,36^. The ATP production declines later, after 18 hpi, when mitochondria are fragmented and mitochondrial DNA is released into the cytosol^27,36,59^. The low cellular level of ATP is one of the factors that lead to apoptosis^60^. The observation of active ATP production at 12 hpi supports the model that apoptosis is activated only at later stages of HSV-1 infection. However, we also show that the cellular proapoptotic processes are supported already at 4 and 8 hpi not only by upregulation of the genes encoding apoptosis-inducing proteins^32,39,40^ but also importantly by downregulation genes encoding apoptosis-blocking proteins^34^. Therefore, the preserved ATP production together with viral antiapoptotic factors retain control over the cellular proapoptotic tools during viral assembly and egress, while this dynamic balance likely shifts toward activation of apoptosis as the infection proceeds further.

Our data reveal that the transcription of mitochondrial genes associated with the mitochondrial organization is extensively regulated during the progression of HSV-1 infection. Specifically, the downregulation of the genes encoding proteins involved in mitochondrial fission could lead to mitochondrial elongation (Fig. 7a). This is supported by a recent soft X-ray microscopy study also showing that HSV-1 infection leads to mitochondrial elongation^28^. Elongation has previously been observed in the dengue virus infection, in which mitochondrial fusion is induced by the inactivation of mitochondrial fission protein, dynamin-related protein 1 (DRP1)^61^. The balance between mitochondrial fusion and fission machinery is changed during the progression of apoptosis leading to mitochondrial fragmentation and blebbing, activation of caspases, and cytochrome c release from mitochondria^62^. The presence of elongated mitochondria in HSV-1-infected cells at 8-12 hpi supports our other findings that the early events of apoptosis including mitochondrial fission have not yet been activated. Notably, our observation of HSV-1 infection-induced relocalization of mitochondria closer to the nuclear envelope in fibroblast (Fig. 7a) is supported by studies in neurons and keratinocytes^27,63^. It has been shown earlier that the cytoplasmic distribution of mitochondria in noninfected cells is regulated by microtubule motor protein contacts with mitochondrial outer membrane protein, mitochondrial Rho GTPase 1 (MIRO1). Deleting MIRO1 causes the repositioning of mitochondria close to the nucleus and results in elevated local concentrations of ATP, H_2_O_2_, and Ca^2+^ in the cytoplasm close to the nucleus^64^. MIRO1 is a calcium sensor protein and elevated cytoplasmic [Ca^2+^] results in a decrease in mitochondrial dynamics or motion standstill in neurons^65,66^. The intracellular [Ca^2+^] increases in HSV-1 infection of neural cells and it leads to a significant reduction of MIRO1-mediated mitochondrial mobility at late stages of infection^67^. The HSV-1 infection-induced elevated level of [Ca^2+^] could also explain the perinuclear distribution of mitochondria in MEF cells. Moreover, the multifunctional HSV-1 ICP34.5 protein which regulates mitochondrial dynamics could have a potential role in the positioning of mitochondria. ICP34.5 interacts with mitochondrial phosphatase PGAM5 which is responsible for the transport and perinuclear localization of mitochondria under infection-induced stress conditions in neural cells^68^.

**Figure 7.**
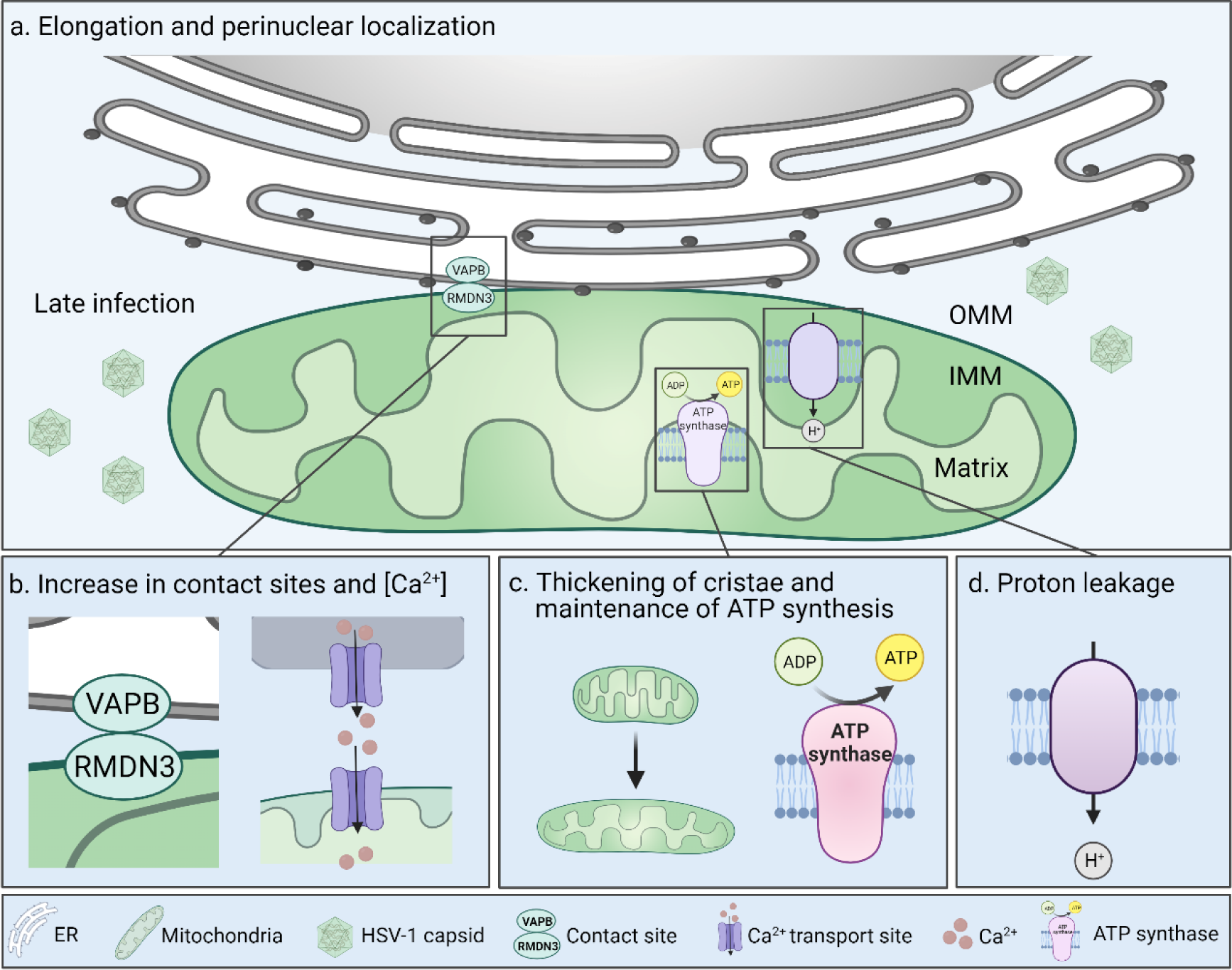
Mitochondrial structure and function are altered in response to progression of herpesvirus infection. **(a)** Progression of HSV-1 infection from 4 to 8-12 hpi triggers mitochondrial elongation and repositioning to the perinuclear region. (**b)** The number of the ER-mitochondria membrane contact sites tethered by VAPB and RMDN3 is increased at late infection and mitochondrial Ca^2+^ content is elevated. (**c**) The progression of infection leads to the thickening of mitochondrial cristae and recovery of ATP production to the level of noninfected cells. (**d**) At the same time, the infection stimulates proton leakage across the mitochondrial inner membrane. VAPB, vesicle-associated membrane protein B; RMDN3, regulator of microtubule dynamics 3; OMM, outer mitochondrial membrane; IMM, inner mitochondrial membrane.

Multiple mitochondrial functions are regulated through the ER-mitochondria membrane contact sites in the outer mitochondrial membrane^69^. Besides being essential for mitochondrial signaling, buffering cytosolic Ca^2+^ level by uptake, division, and metabolism, the contact sites are known to be involved in the infection progress-related balancing of cellular pro- and antiviral responses. Recently Cook et al. elegantly demonstrated that in HCMV, HSV-1, Influenza A, and beta-coronavirus HCoV-OC43 infection the timely recruitment of ER-mitochondria linkers (VAPB and RMDN3) is related to the proceeding of the viral replication^70^. The proteomics analysis also showed that the late HSV-1 infection results in an increased number of other ER-mitochondria contact site proteins, ER ribosome-binding protein 1 (RRBP1), and mitochondrial synaptojanin 2 binding protein (SYNJ2BP)^70,71^. Consistently, our volume EM data, expansion microscopy, and PLA interaction analysis revealed an extensive growth in the number, volume, and density of contact sites at late infection simultaneously with an increase in mitochondrial membrane roughness and mitochondrial Ca^2+^ content (Fig. 7b). The increased availability and clustering of contact sites most likely result in enhanced Ca^2+^ flux from the ER to mitochondria^11,34^ thereby explaining the reactivation of respiration at late infection and possibly also the perinuclear localization of mitochondria. These events are supported by our results showing that the expression of the mitochondrial pore-forming subunit of calcium voltage-dependent calcium channel subunit (CACNA1B) is upregulated in infection.

In HCMV infection the increase in ER-mitochondria contact sites and ER-to-mitochondria transfer of Ca^2+^ are accompanied by cristae remodeling^70,72^. Our studies identified HSV-1 infection-induced upregulation of genes responsible for cristae organization and ATP generation, and downregulation of genes with a negative impact on cristae formation. Our FIB-SEM data also demonstrated that the infection-induced elongated mitochondria contained shorter and thicker lamellar cristae (Fig. 7c). The shortening and widening of the cristae give higher overall curvature to them. Notably, it has been shown that the ATP synthases are preferentially located in the more curved areas, and the energetic efficiency of e.g. heart cells is supported by increased curvature of their cristae^73^. This suggests that the structural alteration of the cristae shape may support the energy generation in infected cells. Consistently, our analysis of the energetic metabolism verified that ATP synthesis was maintained at 8 hpi, (Fig. 7c). Previously published studies showed that the HCMV-induced remodeling of cristae can stimulate cellular respiration^74^. We also showed that the HSV-1 infection is accompanied by increased proton leakage into the mitochondrial matrix independent of ATP synthesis (Fig. 7d). The activation of ATP production and reduced membrane potential seem to have opposite roles in viral replication. However, in dengue 2 virus infection the proton leak was accompanied by an increased ATP generation^75^, and the leakage may also benefit infection as it has been shown to reduce mitochondrial antiviral signaling and response^76^.

Our multimodal integration of advanced imaging, transcriptomics, and metabolomics draws a comprehensive picture of time-dependent changes in mitochondria as HSV-1 infection proceeds from early to late infection. Our results show how the progression of infection shifts the balance from healthy to diseased cells and leads to profound perturbations in mitochondrial homeostasis.

## Data availability

The authors declare that the data supporting the findings of this study are available within the article and its Supplementary Information files, or are available from the corresponding author upon request. GRO-Seq data were deposited into the Gene Expression Omnibus database under accession number GSE243613 and are available at the following URL: xxx. The reviewer access token is inwbyckmnbwfrgi.

## CONTRIBUTIONS

E.M. and M.U.K designed the GRO-seq studies, E.M., S.M. and H.N. performed experiments, and S.L., H.N., S.M. and M.U.K. analyzed data. S.L., V.R., M.H., K.F., T.M., E.P., C.A.L. and M.V.-R. designed cryo-SXT studies, S.L., V.R., V.A., J.H.C. and A.J.P. performed experiments, and S.L., A.A.E., S.K. and V.W. analyzed data. A.G., E.J., and M.V.-R. designed EM, SBF-SEM, and FIB-SEM studies, A.G. and H.V. performed experiments, and S.L., K.K. and I.B. analyzed data. V.R. and S.M. designed the expansion microscopy studies, V.R. performed experiments, and V.R., S.L. and V.A. analyzed data. S.M. designed the PLA studies, S.M. performed experiments, and S.M., S.L. and V.A. analyzed data. S.L., P.T., E.D. and V.H. designed the seahorse studies, S.L. performed experiments, and S.L. analyzed data. M.H., E.D., V.H., E.P., T.M., K.F., M.U.K., E.J. C.A.L. and M.V.-R. administered the experiment, and V.A. and M.V.-R. wrote the manuscript, S.L., V.R., A.G., and S.M. contributed to the results and methods sections, S.M. created Figure 7 with BioRender.com, V.R., E.M., E.D., V.H. and S.M. edited the manuscript. All authors contributed extensively to the work presented in this paper, discussed the results and implications, and commented on the manuscript.

## ETHICS DECLARATIONS

The experiments were done using commercially available cell lines without ethical challenges.

## ELECTRONIC SUPPLEMENTARY MATERIAL

Supplementary material includes methods, supplementary figures, tables, and movies.

## ACKNOWLEDGEMENTS

We thank the staff of the National Center for X-ray Tomography (NCXT) and Advanced Light Source (Lawrence Berkeley National Laboratory, Berkeley, CA), the MISTRAL (BL09) beamline at ALBA Synchrotron (Barcelona, Spain), Diamond Light Source (Oxford, UK), and SiriusXT Limited (Dublin, Ireland) for providing cryo-SXT experiments. We thank Arja Strandell, Mervi Lindman, and Mervi Laanti for technical assistance with EM samples, and the Electron Microscopy Unit (EMBI) of the Institute of Biotechnology, University of Helsinki (Helsinki, Finland) for EM sample preparation and volume EM imaging. We thank the technical support of Sami Salminen, Jänne Kärnä, and Niklas Kähkönen with Seahorse experiments done in BioMediTech (Faculty of Medicine and Health Technology, Tampere University, Tampere, Finland), David Rogers, Stephen O’Connor (SiriusXT) for imaging of cryo-SXT samples, and Niilo Joutsenlahti for GRO-seq sample preparation. We acknowledge Biocenter Finland for infrastructure support.

